# A Molecular Basis of Human Brain Connectivity

**DOI:** 10.1101/2023.07.20.549895

**Authors:** Bernard Ng, Shinya Tasaki, Kelsey M Greathouse, Courtney K Walker, Ada Zhang, Sydney Covitz, Matt Cieslak, Ashley B Adamson, Julia P Andrade, Emily H Poovey, Kendall A Curtis, Hamad M Muhammad, Jakob Seidlitz, Ted Satterthwaite, David A Bennett, Nicholas T Seyfried, Jacob Vogel, Chris Gaiteri, Jeremy H Herskowitz

**Affiliations:** Rush Alzheimer’s Disease Center, Rush University Medical Center; Chicago, Illinois, USA, 60612; Department of Neurology, Center for Neurodegeneration and Experimental Therapeutics, University of Alabama at Birmingham; Birmingham, AL, USA, 35294; Penn/CHOP Lifespan Brain Institute, University of Pennsylvania, Philadelphia, PA, USA; Department of Psychiatry, University of Pennsylvania, Philadelphia, PA, USA; Penn Lifespan Informatics and Neuroimaging Center, University of Pennsylvania, Philadelphia, PA, USA; Department of Child and Adolescent Psychiatry and Behavioral Science, The Children’s Hospital of Philadelphia, Philadelphia, PA, USA; Department of Clinical Science, Malmö, SciLifeLab, Lund University, Lund, Sweden; Department of Biochemistry, Emory University School of Medicine; Atlanta, GA, USA, 30322; Department of Psychiatry, SUNY Upstate Medical University, Syracuse, NY 13210

## Abstract

Neuroimaging is commonly used to infer human brain connectivity, but those measurements are far-removed from the molecular underpinnings at synapses. To uncover the molecular basis of human brain connectivity, we analyzed a unique cohort of 98 individuals who provided neuroimaging and genetic data contemporaneous with dendritic spine morphometric, proteomic, and gene expression data from the superior frontal and inferior temporal gyri. Through cellular contextualization of the molecular data with dendritic spine morphology, we identified hundreds of proteins related to synapses, energy metabolism, and RNA processing that explain between-individual differences in functional connectivity and structural covariation. By integrating data at the genetic, molecular, subcellular, and tissue levels, we bridged the divergent fields of molecular biology and neuroimaging to identify a molecular basis of brain connectivity.

**One-Sentence Summary:** Dendritic spine morphometry and synaptic proteins unite the divergent fields of molecular biology and neuroimaging.

## Main Text

Interactions across biophysical scales, from molecules to tissues, are fundamental to human brain connectivity. Both molecular and neuroimaging studies have independently identified correlates of brain function and cognition, but the search for the molecular basis of functional connectivity is nascent. Recent neuroimaging genetics studies are beginning to address the question of which molecules participate in functional connectivity (*1–5*). However, despite thousands of samples, these studies have found only a dozen independent genetic loci with modest heritability (*6*, *7*). Moreover, most of those genetic variants reside in noncoding regions, making it difficult to determine the implicated genes and mechanisms of action. Another class of studies test for spatial correspondence between gene expression and brain connectivity to infer the implicated genes (*8–12*). This class of studies typically combines the Allen Human Brain Atlas (AHBA) (*13*), which contains gene expression profiles in hundreds of brain regions from six individuals, with functional magnetic resonance imaging (fMRI) data from a different cohort. A limitation to combining data from separate cohorts is that it does not examine the covariation between gene expression and functional connectivity across individuals, but modeling such covariation is important for finding molecules that explain individual variation in functional connectivity and the associated cognitive differences (*14*). Therefore, we gathered data types that span multiple biophysical scales from a cohort of 98 individuals to bridge neuroimaging and molecular biology. Data types included resting state fMRI, structural MRI, and genetic as well as dendritic spine morphometric, proteomic, and gene expression measurements from postmortem tissues of the superior frontal gyrus (SFG) and inferior temporal gyrus (ITG). Based on the stability of functional connectivity patterns within individuals (*15*), we hypothesized that it is possible to combine postmortem molecular and subcellular data with antemortem neuroimaging data to find a molecular basis of brain connectivity. To test this hypothesis, we built models that integrate protein measurements with dendritic spine morphometry in explaining between-individual variation in functional connectivity. As replication, we repeated the analysis with gene expression measurements and structural covariation as a surrogate of connectivity.

To generate an overview of the major molecular systems present in each brain region, we clustered the measured proteins into co-varying sets (modules) in a purely data-driven manner (*16*). Considering synaptic transmission is fundamental to region scale connectivity, we focused our analysis on protein modules (pMod) whose members are enriched for synaptic gene ontology (GO) terms (pMod6 for SFG and pMod8 for ITG in Table S1). We represented the synaptic module of each brain region by the average protein abundance of its members. Accounting for confounding factors including age at death, age of scan, sex, years of education, scanner, post mortem interval, side of brain the molecular data were acquired, and motion, we tested if the synaptic modules of SFG and ITG (their main effects and interaction in aggregate) are associated with an fMRI estimate of their connectivity (Fig 1). This initial test did not detect an association between the synaptic modules and SFG-ITG connectivity (p=0.6839), possibly due to these modules being partly enriched for additional molecular sub-processes. We therefore tested if cellular contextualization of the synaptic modules can better reveal their association with functional connectivity. To do this, we estimated the dendritic spine component of the synaptic modules by fitting the average protein abundance of their members with morphometric attributes of dendritic spines (Fig. S1), and repeated the analysis. Using the dendritic spine component of the synaptic modules indeed resulted in an association with SFG-ITG connectivity (p=0.0174). We next sought to test whether the association is specific to the SFG-ITG connection. Such a test would lend credence to the finding and determine the extent to which connectivity across the brain can be explained by dendritic spines and proteins gathered from SFG and ITG. Accordingly, we associated the dendritic spine component of SFG and ITG synaptic modules with connectivity of all other region pairs. We found only 3% of other connections have higher association strength (Fig. 2A), and these connections generally emanate from either SFG and its nearby areas in the frontal cortex or from ITG. This result likely reflects how spatially proximal areas tend to have similar functional and molecular architectures (*17*, *18*). Moreover, the interaction between the synaptic modules of SFG and ITG explains additional variance beyond their main effects (p=0.0140), with only 1.5% of other connections showing stronger interaction effects. This result suggests that not only is the synaptic module of each region linked to SFG-ITG connectivity, but also the collective molecular state of these regions plays a role.

**Figure 1.**
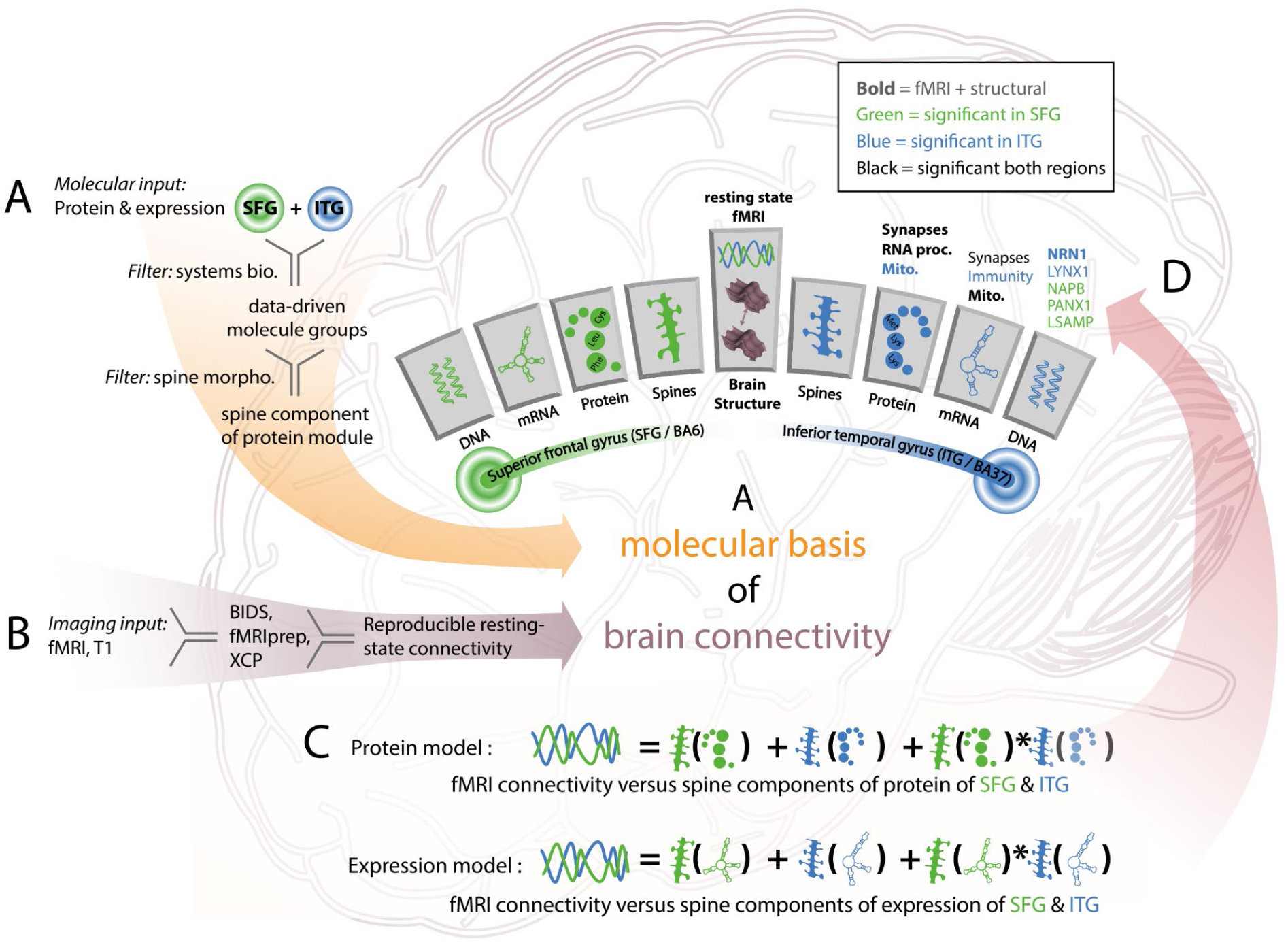
Overview of data types, association models, and key findings. (A) Dendritic spine morphometric, genome-wide gene expression, and protein abundance data were acquired from SFG and ITG in a cohort of 98 individuals. (B) Structural and functional MRI data were acquired from the same 98 individuals. (C) Molecular measurements were integrated with dendritic spine morphologies to model functional connectivity and structural covariation between SFG and ITG. (D) Key molecular processes and proteins associated with SFG-ITG connectivity.

**Figure 2.**
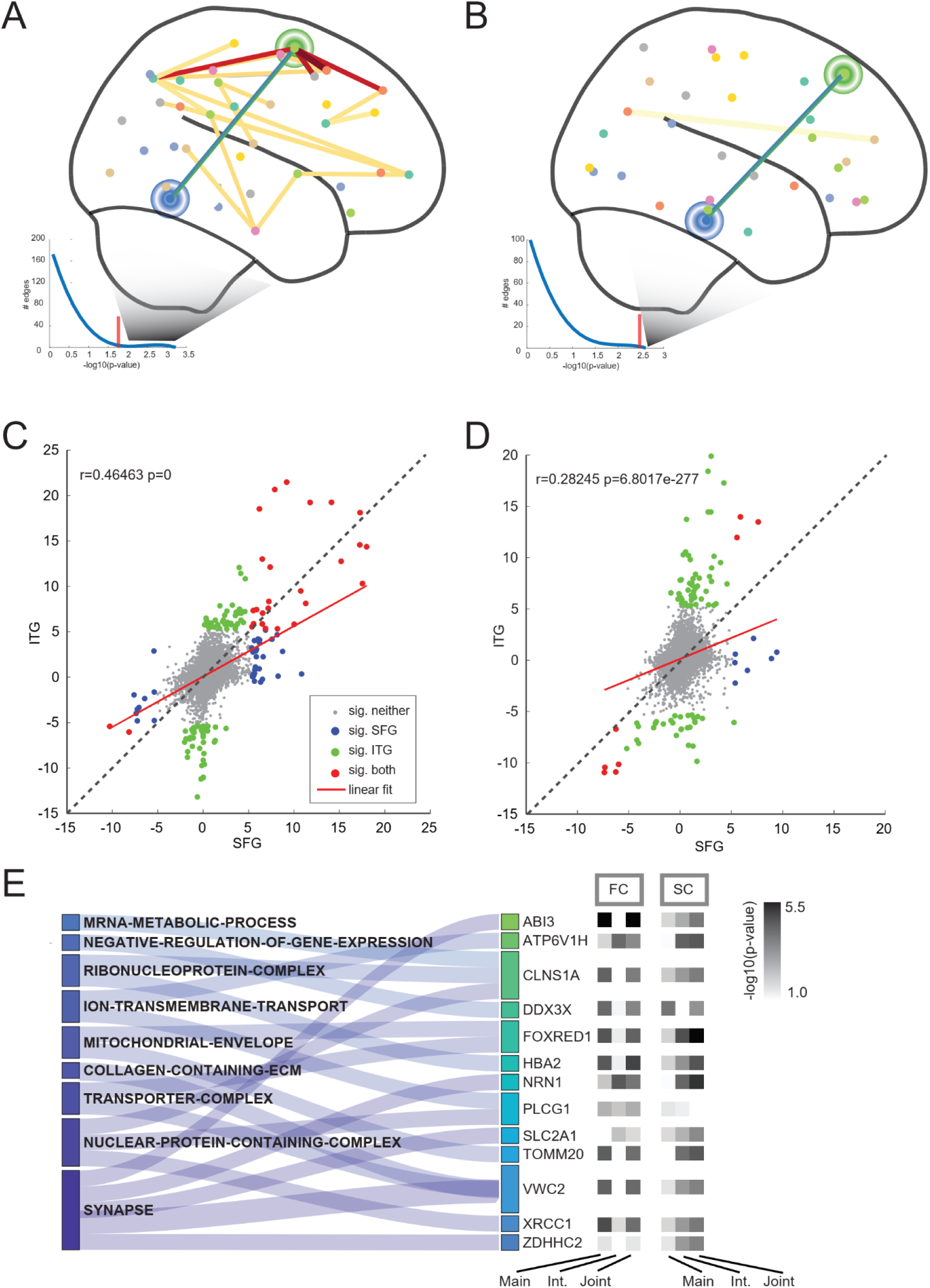
Associations between molecular measures and brain connectivity. (A) SFG-ITG connection and other connections with stronger association strength between functional connectivity and the dendritic spine component of synaptic protein modules displayed. 3% of other connections across the brain showed higher association strength. Connections emanating from SFG or ITG displayed in red. Other connections displayed in yellow. (B) The structural covariation counterpart displayed. 0.4% of other connections showed higher association strength between structural covariation and the dendritic spine component of synaptic protein modules. (C) Comparison of GO enrichment between connectivity-related SFG and ITG proteins. (D) Comparison of GO enrichment between. (E) Functional properties of proteins and their associations with functional connectivity in ITG.

Due to the interplay of functional connectivity and structural morphologies (*19–22*), we repeated the above analysis with an MRI estimate of structural covariation between SFG and ITG as the outcome. Similar to the functional connectivity results, we did not find an association between SFG-ITG structural covariation and the synaptic protein modules (p=0.8815), unless we hone into their dendritic spine component (p=0.0034). We also observed regional specificity, with only 0.4% of other brain region pairs showing stronger association (Fig. 2B). Furthermore, the interaction effect is significant (p=0.0203), with only 3% of other brain region pairs showing stronger effects. The significant associations with the same synaptic protein modules suggest that functional connectivity and structural covariation likely share molecular processes.

We further repeated the analysis with gene expression modules in place of protein abundance modules as replication. The synaptic modules built from gene expression data (eMod5 for SFG and eMod4 for ITG in Table S2) contain a significant number of genes that overlap with members of the synaptic protein modules (OR=2.92, p=2.71×10^-32^ for SFG and OR=2.53, p=5.14×10^-23^ for ITG). Similar to the protein module results, we did not find an association between SFG-ITG connectivity and synaptic expression modules (p=0.0532), unless we focus on their dendritic spine component (p=0.0396). The observed regional specificity with protein abundance modules is also evident with gene expression modules in line with previous transcriptomic studies (*13*, *23*, *24*), with only 2% of other connections displaying stronger association (Fig. S2A). However, we did not observe an interaction effect (p=0.4824). When we associated SFG-ITG structural covariation with synaptic expression modules, we again did not find an association (p=0.9705), unless we hone into their dendritic spine component (p=0.0234). Regional specificity is lower compared to synaptic protein modules, with 14% of other brain region pairs showing stronger association (Fig. S2B), and we did not observe an interaction effect (p= 0.2538). These results show that the expression data replicates the associations between the dendritic spine component of synaptic modules and both functional connectivity and structural covariation, but with weaker interaction effects compared to protein data. More broadly, results above indicate the benefits of cellular contextualization by using dendritic spine morphometry to bridge systems at the molecular level to connectivity at the brain region level and establish their associations.

The associations presented thus far are at the level of modules, which comprise hundreds of proteins and genes. To prioritize specific proteins among the 7788 measured in each region, we modeled SFG-ITG connectivity by the main effect of each protein in a given region, the main effect of the synaptic protein module in the other region, and the interaction between the given protein and the synaptic protein module. We employed this strategy to avoid the limitation on statistical power in testing all pairwise combinations of 7788 proteins between the two brain regions. We prioritized proteins in each brain region based on three effects: the main effect of each protein, its interaction with the synaptic protein module, and these two effects combined. We declared significance at an *α* of 0.05 with false discovery rate (FDR) correction across all proteins, regions, and effects tested. For result interpretation, we applied geneset enrichment analysis (GSEA)(*25*) to the summary statistics of each effect.

Analogous to the module level results, we did not find an association between SFG-ITG connectivity and the abundance of any proteins without cellular contextualization (Table S3). When we repeated the analysis with the dendritic spine component of the proteins, the association strength became stronger (p<10^-4^ for all three effects in both SFG and ITG), and we detected 87 SFG proteins and 344 ITG proteins (Table S3). The SFG proteins are predominantly enriched for GO terms related to synapse, energy metabolism, and RNA processing (Table S4). Top SFG proteins include LSAMP (*26*), NAPB (*27*), KCNIP3 (*28*), and PANX1 (*29*), which are involved in cell surface channel biology at the synapse. Similarly, the ITG proteins are enriched for GO terms related to synapse, energy metabolism, and RNA processing (Table S4). Top ITG proteins include LYNX1, NRN1, PSD2, and RAB11A. Like LSAMP, LYNX1 and NRN1 are glycosylphosphatidylinositol (GPI)-anchored proteins that mediate neuronal receptor activity at excitatory synapses (*30*, *31*). Like NAPB, PSD2 and RAB11A are critical for intracellular signal transduction and protein transport at synapses (*32*, *33*). These findings indicate that while the enriched GO terms of SFG and ITG overlap (OR=48.01, p=1.77×10^-46^, Fig. 2c), their top proteins are different (Fig. S3)

To further probe the relationship between brain function and structure, we repeated the analysis with SFG-ITG structural covariation as the outcome. In contrast to functional connectivity, even without cellular contextualization, we found 72 SFG proteins and 171 ITG proteins associated with SFG-ITG structural covariation (Table S3). Nevertheless, focusing on the dendritic spine component increased association strength (p<10^-4^ for all effects tested except for the main effect of ITG proteins) and detected 625 SFG proteins and 1204 ITG proteins (Table S3). These proteins overlap with proteins detected without cellular contextualization (OR=3.61, p=3.94×10^-5^ for SFG and OR=4.60, p=6.75×10^-20^ for ITG). Interestingly, the enriched GO terms are similar to those found with functional connectivity as the outcome (OR=125.73, p=4.68×10^-26^ for SFG and OR=231.90, p=2.86×10^-98^ for ITG, Table S4), with 18 SFG proteins and 32 ITG proteins in common (Fig. 2E). These findings provide validity for our detected protein-connectivity associations and indicate that certain molecular processes can explain variance in both functional connectivity and structural covariation.

Next, we repeated the analysis with gene expression data, and the overall trend is similar to results obtained with the protein abundance data. We did not find any genes associated with SFG-ITG connectivity without cellular contextualization (Table S5). When we focused on the dendritic spine component of gene expression, the association strength became higher (p<10^-4^ for all tested effects except for the main effect of SFG genes), and we identified 1 SFG gene and 324 ITG genes. The enriched GO terms are predominantly related to synapses, energy metabolism, and RNA processing (Table S6) similar to those found with protein abundance (OR=29.67, p=1.58×10^-42^ for SFG and OR=6.61, p=2.56×10^-20^ for ITG). However, the ranking of genes by association strength does not align with that of proteins (Fig. S4). When we used structural covariation as the outcome, we found association with 130 SFG genes and 18 ITG genes without cellular contextualization. Nevertheless, we detected more genes (1023 SFG genes and 248 ITG genes) and observed higher association strength (p<10^-4^ for all tested effects except for the main effect of SFG genes with p=0.0178) when we focused on the dendritic spine component of gene expression. The enriched GO terms found based on gene expression overlap with those found based on protein abundance (OR=66.57, p=7.29×10^-12^ for SFG and OR=164.97, p=2.66×10^-26^ for ITG), but the ranking of genes and proteins do not align (Fig. S5). Functional connectivity and structural covariation share common GO terms (OR=68.80, p=1.46×10^-45^ for SFG and OR=13.18, p=6.91×10^-10^ for ITG), but have no common genes (Fig. S6). Overall, our results show consistency between protein abundance and gene expression at the molecular function level, even though the proteins and genes involved do not overlap, which is likely due to the low correlation between mRNA level and protein abundance of the same gene (*34*).

To place our connectivity-associated proteins and genes in context with other imaging transcriptomic studies, we compared with genes previously found by combining the AHBA with fMRI data of other cohorts (*35*). The 125 AHBA connectivity-related genes do not place highly among our proteins ranked by their effects on connectivity (Fig. S7), possibly due to post transcriptional regulation, which limits the correlation between mRNA level and protein abundance of the same gene (*34*). Those genes are only enriched among ITG genes ranked by their interaction effects (p=0.01 based on GSEA). This result suggests that the generic set of genes found by correlating spatial patterns of gene expression in AHBA with functional connectivity estimated from other cohorts have only mild correspondence with genes found by modeling covariation between gene expression and SFG-ITG connectivity within the same set of individuals.

Finally, we compared the connectivity-associated proteins and genes found herein with prior imaging genetics studies. The largest related GWAS (*6*) found 4 independent loci (*EPHA3*, *DPP4*, *FBXO11*, and *ZNF326*) that are associated with SFG-ITG connectivity (rfMRI connectivity ICA100 edge 849). These loci were derived by spatially mapping genome-wide significant genetic variants to their closest genes. Among the four loci, *EPHA3* shows the highest heritability in the edge 849 GWAS and the dendritic spine component of *EPHA3* expression displays nominal association with SFG-ITG connectivity in our data (p=0.0192 for SFG main effect and p=0.0140 for ITG main effect). We could not perform the same analysis for protein abundance because tryptic peptides from *EPHA3, DPP4*, *FBXO11*, and *ZNF326* were not measured in our dataset. Instead, we assessed our found proteins in relation to the edge 849 GWAS through a functional mapping approach. For each brain region, we built protein prediction models to combine the effects of genetic variants that are proximal to each protein (*36*). We then applied these models to the summary statistics of the edge 849 GWAS to test whether the genetic component of the proteins is associated with edge 849. Overall, the genetic component of neither SFG proteins nor ITG proteins is associated with edge 849 after FDR correction (Table S7), likely due to limited heritability in edge 849. Accordingly, we focused on the genetic effects of our found proteins. For SFG, 3339 proteins have adequate heritability for model construction. Among these 3339 proteins, the dendritic spine component of 33 proteins are associated with SFG-ITG connectivity in our data (FDR-corrected p<0.05). Among those proteins, the genetic component of 2 proteins, NGB (p=0.0219) and POLR2J (p=0.0423), display nominal associations with edge 849. For the ITG region, 3445 proteins have adequate heritability for model construction. Among these 3445 proteins, the dendritic spine component of 147 proteins are associated with SFG-ITG connectivity in our data (FDR-corrected p<0.05). Among those proteins, the genetic component of 11 proteins, including NRN1 (p=0.0456), display nominal associations with edge 849. These results provide further support for some of the protein-connectivity associations found in our data. The repeated detection of NRN1 (*37*), or neuritin, at the DNA and protein level suggests that this secreted neuropeptide plays a critical role in synaptic biology in supporting brain connectivity of ITG.

The vision for integrating postmortem omics data and antemortem neuroimaging data is to motivate molecular biology experiments that recapitulate the processes underlying connectivity changes in health and disease. Progress toward this vision appears possible, as we found the conjunction of functional connectivity, protein, and cell structure to have numerous associations. A limitation to our study relates to the regional specificity of the findings, as our found genes and proteins are prioritized for connectivity with SFG, ITG, and their nearby brain areas. Extending the brain coverage will require measuring protein abundance in other regions. By focusing major perspectives in neuroscience on the same set of brains, this study lays the groundwork for understanding how human brain function is supported by multiple biophysical scales.

## Acknowledgements

We are grateful to the participants in the Religious Order Study, the Rush Memory and Aging Project (ROSMAP).

## Author contributions

Conceptualization: BN, ST, CG, JH

Methodology: BN, CG, ST, JH

Investigation: BN, ST, CG, JH, TS, JV, KG, CK, SC, MC, AA, JA, EP, KC, HM

Visualization: BN, ST, CG

Funding acquisition: DB, CG, TS, JH

Writing – original draft: BN, CG, ST, JH

Writing – review & editing: BN, ST, AZ, JS, TS DB, NS, JV, CG, JH

## Data and materials availability

Data available through radc.rush.edu and https://adknowledgeportal.synapse.org/

## Materials and Methods

### Cohort

98 subjects of the Religious Orders Study and Rush Memory and Aging Project (ROSMAP) (*38*) have all data types needed for this work, namely functional magnetic resonance imaging (fMRI), structural MRI, genetic, dendritic spine morphometry, RNAseq, and proteomic data. All enrolled participants agreed to annual clinical evaluation and brain donation at death. Both studies were approved by an Institutional Review Board of Rush University Medical Center. Subjects signed informed consent, an Anatomical Gift Act, and a repository consent to allow their resources to be shared. The dendritic spine morphometry, RNAseq, and proteomic data were measured from postmortem tissue samples of the superior frontal gyrus (SFG) and inferior temporal gyrus (ITG). For each subject, tissue samples were drawn from either the left or the right hemisphere but not both sides. 52% of subjects had tissue samples drawn from the left hemisphere. The average age of subjects at time of MRI scan and at death are 88±6 years and 91±6 years, respectively, with an average time interval between MRI scan and age at death of 3±2 years. Average postmortem interval is 8.5±4.6 hours. 77% of the subjects are female, and the subjects have 15±3 years of education. Further demographic details on the subjects were previously described (*38*).

### Proteomic

Multiplex tandem mass tag mass spectrometry (TMT-MS) was employed to generate proteomic data (7788 proteins). Brain tissue homogenization, protein digestion, TMT peptide labeling, high-pH offline fractionation, and the liquid chromatography-tandem mass spectrometry as well as protein quantification, batch correction, and data pre-processing were performed as previously described (*39–41*). Briefly, paired human tissue samples from SFG and ITG were obtained from 98 ROSMAP participants. Approximately 100 mg of tissue was homogenized in 8 M urea, 10 mM Tris, 100 mM NaH_2_PO_4_, pH 8.5 buffer with HALT protease and phosphatase inhibitor cocktail (Thermo Fisher Scientific) using a Bullet Blender (Next Advance). Protein concentration was determined by bicinchoninic acid assay (Pierce) and one-dimensional SDS-PAGE gels were run with Coomassie blue staining for quality control prior to protein digestion. Lysyl endopeptidase (Wako) at 1:100 (wt/wt) was added to each 100 μg sample for digestion overnight. Trypsin (Promega) was added at 1:50 (wt/wt) and digestion occurred for another 16 h. The peptide solutions were acidified and de-salted. The samples were then loaded onto the column, washed, and eluted. An equal amount of peptide from each sample was aliquoted and pooled as the global internal standard (GIS), which was split and labeled in each TMT batch. The eluates were then dried to completeness using a SpeedVac. Peptides from each individual case and the GIS pooled standard or bridging sample were labeled using the TMT 11-plex kit (ThermoFisher 90406). For each batch, up to two TMT channels were used to label GIS standards while the remaining TMT channels were used for samples after randomization. Next, high-pH offline fractionation was performed, and a 96 individual equal volume fractions were collected across the gradient and then pooled by concatenation into 24 fractions and finally dried with a SpeedVac. Fractions were resuspended in an equal volume of loading buffer and analyzed by liquid chromatography-tandem mass spectrometry (LC–MS/MS). Peptide eluents were separated on a self-packed C18 (1.9 μm, Dr. Maisch) fused silica column (25 cm × 75 μM internal diameter, New Objective) by a Dionex UltiMate 3000 RSLCnano liquid chromatography system (Thermo Fisher Scientific). Peptides were monitored on an Orbitrap Fusion mass spectrometer (Thermo Fisher Scientific). For database searching and protein quantification, raw MS data files were analyzed in the Proteome Discover software suite (version 2.3, ThermoFisher) and MS/MS spectra were searched against the UniProtKB human proteome database. Following spectral alignment, peptides were assembled into proteins and filtered based on the combined probabilities of their constituent peptides to a final FDR of 1%. Multi-consensus was performed to achieve parsimony across individual batches. In cases of redundancy, shared peptides were assigned to the protein sequence in adherence with the principles of parsimony. Only peptide spectral matches with less than 50% isolation interference were used for quantification and only unique and razor peptides were considered for quantification. Finally, batch correction and data pre-processing was performed. A total of 10,426 high confidence, master proteins were identified across the 26 TMT batches, but only proteins quantified in >50% of samples were included in subsequent analyses. Log2 abundances were normalized as a ratio dividing by the central tendency of pooled standards. Batch correction was performed using a Tunable Approach for Median Polish of Ratio (https://github.com/edammer/TAMPOR). Multidimensional scaling plots were used to visualize batch contributions to variation before and after batch correction. Network connectivity was used to remove outliers, defined as samples that were greater than 3 standard deviations away from the mean.

### Dendritic Spine Imaging and Processing

Golgi-Cox staining, using the FD Rapid Golgi Stain Kit (FD Neurotechnologies, Catalog #PK401), was employed to visualize dendrites and dendritic spines in postmortem samples from SFG and ITG. The flowing adjustments were made from the standard operating procedure in the manual from the kit. All steps were performed at room temperature. Solutions A (potassium dichromate and mercuric chloride) and B (potassium chromate) were combined 48 hours prior to tissue submersion. 2mL of solution A+B were placed in wells of a 12 well plate (Fisher Scientific, Catalog #08-772-29). Frozen tissue blocks, approximately 10mm x 10mm x 10mm, were dropped into the A+B solution in each well. The chromate solution was replaced after 24 hours and the tissue blocks remained in the solution for 21 days. Next, the tissue blocks were transferred to a new 12 well dish containing solution C. Solution C was replaced after 24 hours. After 72 total hours in solution C, each tissue block was sliced into 125 µm sections in solution C using a Leica Vibratome (VT1000 S). Free floating sections were placed in a 6-well dish (Fisher Scientific, Catalog #353046) that contained solution C in each well. Tissue sections were sequentially moved from solution D, to solution E, and to distilled water similar to manufacturer’s instructions. Slices were dehydrated with alcohols (70%, 90%, 100% Ethanol) and cleared with xylenes (Fisher Scientific, Catalog #X3P). Slices were then placed on glass slides (Fisher Scientific, Catalog #12-550-15) with a single slide spacer (Electron Microscopy Sciences, Catalog #70327-20S), sealed with Permount (Fisher Scientific, Catalog #SP15-100), and coverslipped with 24 x 50mm, thickness 0.13-0.17mm, glass (Carolina Biological, Catalog #633153). Slides were dried in the dark for 7 days prior to microscopy imaging.

Dendrites were imaged by blinded experimenters using brightfield microscopy. Each case had multiple slides available for imaging. From each tissue section, 1-2 pyramidal neurons were imaged for analysis. Between 8-12 neurons were sampled per individual. A single dendritic segment was imaged per neuron. Dendrite segments were imaged if the following criteria were met; (1) located centrally within the total tissue depth, (2) not obscured by staining debris and/or intersecting neighboring neurons, (3) fully impregnated. Additionally, only dendrites that were over 30 µm in length and had an approximate diameter of 1 µm were imaged. Tissue slices were visualized under 10X magnification to determine tissue quality. Once suitable dendrites were identified, tissue slices were visualized at 60X magnification using Type F immersion oil (Nikon, Catalog #MXA22168). Z stacks were captured at a step size of 0.1 µm at 60X using a Nikon Plan Apo 60X/1.40 numerical aperture (NA) oil-immersion objective in combination with a high-NA oil condenser (Nikon, Catalog #MEL59500) on a Nikon Eclipse Ti2 inverted microscope with a Lumencor SOLA light engine and Hamamatsu ORCA-flash 4.0 digital camera. All images were 1024 x 1024 pixels.

Reconstructions of dendrites and dendritic spines were conducted by blinded experimenters using Neurolucida 360 (MBF Biosciences, version 2.70.1). Nikon ND2 files were converted to 16-bit TIFF files using ImageJ, then imported to Neurolucida 360. Dendritic segments were traced using a user-guided semi-automated directional kernel algorithm. Initiation and termination points for dendrite reconstruction were determined with the following criteria: (1) ≥ 5 µm away from the distal tip of the dendrite, (2) consistent diameter, (3) level axis with limited smear in the z plane, and (4) ≥ 20 µm in length. After the dendritic segment was traced, the experimenter verified that the points located on the dendrite were accurate in X, Y, and Z planes, and if not, made manual adjustments to the trace. The diameter of the dendrite at each point was also verified to ensure accuracy of the trace. Dendritic spines were traced using voxel clustering. The following parameters for used for spine identification: outer range, 7 µm; minimum height, 0.3 µm; detector sensitivity, 90 to 125%; minimum count, 8 voxels. The morphology of each dendritic spine was examined to verify that axial smear did not cause misrepresentation. Merge and slice tools in Neurolucida 360 were employed to correct inaccuracies. The position of every spine’s backbone point was verified by the blinded experimenter. If the backbone caused a misrepresentation, the dendritic spine was visualized from the Z-plane, and the experimenters re-oriented backbone points in the X-Y plane. Notably, repositioning in the X-Z or Y-Z plane was performed while the spine was being viewed from the lateral angle. Morphometric analysis was conducted for each spine, and measurements categorized spines into thin, stubby, mushroom, and filopodia classes. The following parameters were used for spine classifications: head-to-neck ratio, 1.1; length-to-head ratio, 2.5; mushroom head size, 0.35 µm; filopodium length, 3.0 µm. Spines with a head-to-neck ratio > 1.1 and head diameter > 0.35 µm were classified as mushroom. Spines were classified as filopodia or thin if head-to-neck ratio was < 1.1 and either (1) length-to-head ratio was > 2.5 or (2) head size was < 0.35 µm. Moreover, if the total length of the spine was > 3.0 µm, the spine was classified as filopodia, and if the total length was < 3.0 µm, then the spine was classified as thin. Dendrite and spine reconstructions were exported to Neurolucida Explorer (MBF Biosciences, version 2.70.1), where data were collected for quantitative analysis. The dendritic spine measurements were exported and collected in Microsoft Excel. Spine density was calculated by determining the quantity of spines per 10 µm of dendrite length. Spine length was defined as the curvilinear backbone length from the insertion point to the most distal point of the spine head. Head diameter was defined as the breadth of the spine head at its widest cross-sectional point.

### Gene Expression

RNA was extracted using the Chemagic RNA tissue kit (Perkin Elmer, CMG-1212). RNA was concentrated (Zymo, R1080), and RQN (RIN score) was calculated using Fragment Analyzer (Agilent, DNF-471). RNA concentration was determined using the Qubit broad range RNA assay (Invitrogen, Q10211) according to the manufacturer’s instructions. 500ng total RNA was used for RNA-Seq library generation, and rRNA was depleted with RiboGold (Illumina, 20020599). A Zephyr G3 NGS workstation (Perkin Elmer) was utilized to generate TruSeq stranded sequencing libraries (Illumina, 20020599) with custom unique dual indexes (IDT) according to the manufacturer’s instructions with the following modifications. RNA was fragmented for 4 minutes at 85°C. The first strand synthesis was extended to 50 minutes. Size selection post adapter ligation was modified to select larger fragments. According to the manufacturer’s instructions, library size and concentrations were determined using an NGS fragment assay (Agilent, DNF-473) and Qubit ds DNA assay (Invitrogen). The modified protocol yielded libraries with an average insert size of around 330-370bp. Libraries were normalized for molarity and sequenced on a NovaSeq 6000 (Illumina) at 80-100 million reads, 2×150bp paired-end. RNA-Seq data processing was implemented using three parallel pipelines, an RNA-seq QC pipeline, a gene/transcripts quantification pipeline, and a 3’ UTR quantification pipeline. In the QC pipeline, paired-end RNA-Seq data were first aligned by STAR v2.6 to a human reference genome. The primary assembly of reference genome fasta file and transcriptome annotation came from Gencode (Release 27 GRCh38). Picard tools were applied to the aligned bam files to assess the quality of RNA-Seq data. In the quantification pipeline, transcript raw counts were calculated by Kallisto (v0.46).

We preprocessed the RNAseq data following the pipeline previously described (*42*). In brief, we applied TMM normalization (using edgeR calcNormFactors) to the raw counts to estimate the effective library size of each subject. We then applied voom/limma to regress out confounds and convert the counts into log2(CPM). Technical confounds included batch, study (ROS or MAP), RNA integrity number, postmortem interval, library size, percentages of aligned reads, coding and intergenic bases, ribosomal and untranslated region bases, duplicated reads, 3’ prime bias, 5’ over 3’ prime bias, and coefficient of variation in coverage. Biological confounds, including age, sex, and top 10 expression principal components, were subsequently regressed out. Only genes with mean log2(CPM) > 2 were kept.

### Neuroimaging

Structural scans were acquired using T1-weighted MRI (1.5T GE: 1 mm^3^ resolution, TR = 6.3 ms, TE = 2.8 ms, flip angle = 8°; 3T Siemens: 1 mm^3^ resolution, TR=2300 ms, TE = 2.98 ms, flip angle = 9°; 3T Philips: 1 mm^3^ resolution, TR = 8.0 ms, TE = 3.7 ms, flip angle = 8°). Further details on scanner protocols can be found at https://www.radc.rush.edu/docs/var/scannerProtocols.htm (*43*). Non-uniformity correction, skull-stripping, spatial normalization to the MNI152 template, and tissue segmentation were performed using Advanced Normalization Tools (ANTs). Freesurfer’s “recon-all” function was applied to extract morphometric attributes.

Resting state fMRI BOLD data were acquired on multiple scanners: 1.5T GE Signa scanner (5 mm^3^ resolution, TR/TE=2000/33 ms, flip angle 85°), 3T Siemens Magnetom TrioTim syngo (3.3 mm^3^ resolution, TR/TE=3000/30 ms, flip angle 80°) and 3T Philips Achieva Quasar TX (3.3 mm^3^ resolution, TR/TE=3000/30 ms, flip angle 80°). Further details on scanner protocols can be found at https://www.radc.rush.edu/docs/var/scannerProtocols.htm (*44*). Raw fMRI data were preprocessed using a well validated pipeline of robust and reproducible fMRI processing tools. First, CuBIDS was used to identify and investigate scans with deviant parameters(*45*). fmriprep (v. 20.2.3)(*46*) was then applied to realign and slice-time correct the fMRI volumes, with distortion correction further performed if accompanying fieldmap volumes were available. The resulting fMRI volumes were co-registered to their corresponding T1 volumes using FLIRT. Finally, the fMRI timeseries underwent confound regression using the eXtensible Connectivity Pipeline (XCP; https://github.com/PennLINC/xcp_d)(47) (v. 0.0.4 “xcp-abcd”) with the 36p+Despike model(*48*). In brief, bandpass filtering between 0.01 and 0.08 Hz was applied, followed by despiking with AFNI(*49*) and regression of 36 parameters from the timeseries(*50*). Regressors included six motion attributes derived during realignment, global signal, and two physiological parameters, as well as their derivatives, squares of derivatives, and quadratic terms. Additionally, mean framewise displacement was calculated for each subject and used to account for any residual effect of head motions in subsequent analysis.

### Genetic

Genotyping was performed using the Infinium Global Screening Array, Affymetrix Genome-Wide HumanSNP Array 6.0, and Illumina OmniQuad Express platform (*51*, *52*). The genotyping data underwent sample-level exclusions, including removing duplicated samples, those with a genotyping success rate below 95%, and samples with discordant gender information. At the probe level, additional filtering was applied based on specific quality-control criteria: a Hardy-Weinberg equilibrium p-value threshold (< 1e-50), genotype call rate (< 0.9), and misshape test (< 1e-9). For imputation on the Haplotype Reference Consortium r1.1, we utilized the Michigan Imputation Server, Minimac3 version 1.0.4, and Eagle version 2.3. Before imputation, the input data was prepared using the HRC-1000G-check-bim_ts.pl script, available at https://www.well.ox.ac.uk/~wrayner/tools/. Imputed genotypes with an information score greater than 0.3 were converted to a Plink binary file format using Plink 1.9 for downstream analysis.

### Molecular Module Estimation

To determine the molecular systems of interacting proteins in each brain region, we clustered the proteins into co-abundance modules using a fully data-driven approach called SpeakEasy (*16*), which takes in a similarity matrix and joins nodes into modules based on both local connectivity and global network structure. To estimate the degree of interaction between each pair of proteins, we correlated their abundance levels across subjects. We repeated this estimation for all protein pairs to generate a protein correlation matrix for each region and applied SpeakEasy, which extracted 10 modules for SFG and 10 modules for ITG (Table S1). Considering our goal is to model brain connectivity, which arises from synaptic processes, we applied geneset enrichment analysis (GSEA) (*25*) and focused our analysis on modules (pMod) whose protein members are most enriched for synaptic GO terms. Using this criterion, we selected pMod6 for SFG and pMod8 for ITG, which happen to have the greatest protein overlap among all protein module pairs (OR=13.16, p= 3.99×10^-186^). To represent a protein module, we used the average abundance levels of its protein members. We applied the same procedures to the gene expression data, which extracted 9 modules for SFG and 9 modules for ITG (Table S2). Based on synaptic GO term enrichment, we selected and focused our analysis on eMod5 for SFG and eMod4 for ITG, which also happen to have the greatest gene overlap among all expression module pairs (OR=57.25, p= 4.94×10^-323^).

### Functional Connectivity Estimation

To estimate functional connectivity, we first used a functional atlas comprising 100 parcels (Schaefer2018_100Parcels_17Networks_order_FSLMNI152) generated from resting state fMRI data of >1000 subjects(*53*) to divide the brain into functionally homogenous regions. We then averaged the time-series of voxels within each parcel and computed the Pearson’s correlation between all pairs of parcels for each subject. To select the brain connection (i.e. the parcel pair) most relevant to our molecular data, we examined the spatial overlap between each functional parcel and the brain areas at which we drew tissue samples. Parcels #25 (left hemisphere) and #74 (right hemisphere) have the greatest overlap with our SFG tissue samples, and parcels #14 (left hemisphere) and #65 (right hemisphere) have the greatest overlap with our ITG tissue samples. For associating with the molecular data, we only used the SFG-ITG connectivity estimate that matches the side of brain from which we drew tissue samples for each subject, i.e. connectivity estimate between parcels #25 and #14 if we drew brain tissue samples from the left hemisphere for a given subject, and vice versa. We used the connectivity estimates of other parcel pairs for assessing regional specificity (see Module Level Association Analysis below).

### Structural Covariation Estimation

To estimate structural covariation, we used the 9 regional morphological attributes (number of vertices, surface area, gray matter volume, average cortical thickness, standard deviation of cortical thickness, mean curvature, Gaussian curvature, fold index, and curvature index) from Freesurfer cortical surface reconstruction with regions defined anatomically based on the DKT atlas (*54*). Let **Z***_i_* be the *n*×9 morphological attribute matrix of region *i* with *n* being the number of subjects. The standard approach (*55*) estimates the structural covariation between regions *i* and *j* by correlating row *k* of **Z***_i_* with row *k* of **Z***_j_* for *k* = 1 to *n*. The assumption is that morphological attributes have a one-to-one correspondence across regions, i.e. fold index in region *i* co-varies with fold index in region *j*, curvature index in region *i* co-varies with curvature index in region *j*, etc. However, fold index of region *i* might also co-vary with e.g. curvature index of region *j*. Therefore, to mitigate the one-to-one attribute correspondence assumption, we applied canonical correlation analysis (CCA) to find combinations of morphological attributes that maximally co-vary between region pairs. Specifically, we applied CCA to **Z***_i_* and **Z***_j_* to find 9×9 projection matrices **A***_i_* and **A***_j_* that maximizes their correlation, i.e. max *tr*(**A***i*^T^**Z***i*^T^**Z***_j_***A***_j_*) s.t. **A***i*^T^**Z***i*^T^**Z***_i_***A***_i_* = **I** and **A***j*^T^**Z***j*^T^**Z***_j_***A***_j_* = **I**. Adopting a recent approach where co-fluctuation in fMRI intensity at single time point resolution is used to estimate connectivity between two regions (*56*), we used **S***_ij_* = (**Z***_i_***A***_i_*) ◦ (**Z***_j_***A***_j_*) as estimates of structural covariation between regions *i* and *j*, where ◦ denotes element-wise product and each CCA dimension (i.e. each column of **S***_ij_*) captures a different mode of shape co-variation. For associating with the molecular data, we split the subjects based on the side of brain from which we drew tissue samples, and separately applied CCA to those two sets of subjects. We focused our analysis on the structural covariation between “superiorfrontal” and “inferiortemporal” in the DKT atlas, which best overlap with where we sampled brain tissues, and used other anatomical region pairs for assessing regional specificity (see Module Level Association Analysis).

### Module Level Association Analysis

Let **Y***_ij_* be a *n*×1 vector corresponding to neuroimaging estimates of a brain attribute (functional connectivity or structural covariation between regions *i* and *j*) of *n* subjects, **X***_i_* and **X***_j_* be *n*×1 vectors corresponding to the average molecular values (protein abundance or gene expression level) of members within the synaptic modules of regions *i* and *j*, and **C** be a *n*×*d* confound matrix. We modeled **Y***_ij_* as: **Y***_ij_* = *α*_0_ + ***Cα*** + **X***_i_β_i_* + **X***_j_β_j_* + **X***_i_*◦**X***_j_β_ij_* + ɛ, with age at death, age of scan, sex, years of education, scanner, post mortem interval, side of brain molecular data were acquired, and mean framewise displacement as confounds. Our primary question is whether **X***_i_β_i_* + **X***_j_β_j_* + **X***_i_*◦**X***_j_β_ij_* can significantly explain variance in **Y***_ij_* across subjects for *i* = SFG and *j* = ITG. Due to the synaptic modules being partly enriched for other molecular subprocesses, we tested whether cellular contextualization of these modules can better reveal their association with our brain attributes of interest. Accordingly, we extracted the dendritic spine component of the synaptic modules by fitting the average molecular values of their members with morphometric attributes of dendritic spines, and used these components as **X***_i_* and **X***_j_*. To assess regional specificity, we fitted the same model with **Y***_ij_* set to brain attribute values of other region pairs (i.e. non SFG-ITG connections) and computed the percentage of other region pairs with lower p-values. For associating with structural covariation, which has 9 CCA dimensions, we separately fitted the model to each CCA dimension and applied aggregated Cauchy association test (ACAT) to combine p-values across the 9 dimensions (*57*). We opted to use ACAT over other p-value combination tests because it provably provides optimal power in sparse settings (*57*, *58*), which accentuates the CCA dimensions that have stronger association with the synaptic modules, in contrast to e.g. Fisher’s method that weights all dimensions equally.

### Molecule Level Association Analysis

To find the specific molecules (proteins and genes) associated with our brain attributes of interest, we modeled **Y***_ij_* as: **Y***_ij_* = *α*_0_ + ***Cα*** + **x***_mi_β_mi_* + **X***_j_β_j_* + **x***_mi_*◦**X***_j_β_mij_* + ɛ, where **x***_mi_* is a *n*×1 vector of molecular values (abundance level of protein *m* or expression level of gene *m* in region *i*) of *n* subjects. The other variables are as defined in the module level association model. We used this model to avoid testing all pairwise combinations of molecules between SFG and ITG (7788^2^ protein pairs or ∼12,000^2^ gene pairs), which limits statistical power. We tested for molecules involved in SFG by setting *i* to SFG and *j* to ITG, and vice versa. We focused on testing 3 terms: **x***_mi_β_mi_*, **x***_mi_*◦**X***_j_β_mij_*, and **x***_mi_β_mi_* + **x***_mi_*◦**X***_j_β_mij_*, i.e. effects that do not exclusively involve the synaptic modules. We declared significance at 0.05 with false discovery rate (FDR) correction over the combinations of molecules, regions, and effects of interests. We considered a molecule as significant if any of the 3 effects is significant with FDR correction. Considering a molecule could be involved in multiple molecular processes, we again applied cellular contextualization by substituting the molecular values by their dendritic spine components and repeated the analysis. To test if cellular contextualization improves association strength, for each combination of effect and region, e.g. main effect of SFG, we compared the –*log*_10_(*p*) of an effect with vs. without cellular contextualization using a permutation test. Let **d** be a *q*×1 vector with elements corresponding to the differences in –*log*_10_(*p*) between with and without cellular contextualization for *q* molecules. For each permutation, we randomly selected *q*/2 molecules, flipped the sign of their –*log*_10_(*p*) differences, and computed the average difference over molecules. We repeated this procedure 10^4^ times and computed the proportion of times the non-permuted average –*log*_10_(*p*) difference is smaller than the permuted average *-log*_10_(*p*) differences, i.e. a permutation-based p-value. We further performed GSEA (*25*) on GO terms and a connectivity-related geneset (*35*) for result interpretation. We applied GSEA on t-values of all measured molecules (separately for proteins and genes) for each of the 3 effects on functional connectivity (separately for SFG and ITG) and declared significance at 0.05 with Bonferroni correction (p<5×10^-6^). We considered a GO term as significant if any of the 3 effects is significant. For **x***_mi_β_mi_* + **x***_mi_*◦**X***_j_β_mij_*, we only have a F-value for each molecule. Therefore, we approximated a t-value for each molecule by taking the square root of its F-value and multiplying it by the sign of the mean t-value of **x***_mi_β_mi_* and **x***_mi_*◦**X***_j_β_mij_*, i.e. take the sign of the more dominant effect. For structural covariation, we only have p-values derived by aggregating the CCA dimensions. Therefore, for each molecule, we approximated its effect direction by averaging its t-values on a given effect across the CCA dimensions and taking the sign of this average. We used signed –*log_10_*(p) as input to GSEA.

### MetaXcan Analysis

To compare our connectivity-associated proteins against prior imaging genetics studies, we used a functional mapping approach, called MetaXcan (*36*), to convert SNP level GWAS summary statistics to protein level. For each protein *m*, We first built a model to extract the genetic component of its protein abundance: **P***_m_* = Σ*_sɛSm_* **w***_ms_***S***_s_* + ***ε****_m_*, where **P***_m_* is a *n*×1 vector containing the abundance levels of protein *m*, **S***_s_* is a *n*×1 vector containing the dosage of SNP *s*, **w***_ms_* is the *s*^th^ element of a *l_m_*×1 model weight vector, **w***_m_*, to be estimated, and *S_m_* is the set of *l_m_* SNPs within ±1Mb from the transcription start site of gene *m* that produces protein *m*. Following MetaXcan, we estimated **w***_ms_* using elastic net regression by applying GLMNET with its default settings, i.e. 10 fold cross-validation to set the sparsity parameter. We regressed out confounds including age of death, sex, post mortem interval, top 3 ancestry principal components, and top 10 protein principal components from the protein abundance measurements prior to fitting the models. Given **w***_ms_*, protein level GWAS z-score, **z***_m_*, of protein *m* can be estimated by: **z***_m_* = Σ*_sɛSm_* **w***_ms_**σ**_s_*/***σ****_m_ ·* **z***_s_*, where **z***_s_* is the GWAS z-score of SNP *s*, and ***σ****_m_* and ***σ****_s_* are the variance of protein *m* and SNP *s*, respectively. We built models for SFG and ITG proteins separately, and packaged the models into SQLITE databases (https://doi.org/10.7303/syn52087434) so users can directly use them with the public MetaXcan software (https://github.com/hakyimlab/MetaXcan). We applied MetaXcan with our protein models to a GWAS on functional connectivity between SFG and ITG (rfMRI connectivity ICA100 edge 849 (*6*)), and examined which proteins found associated with SFG-ITG connectivity in our data show significant genetic effects based on **z***_m_*.

## Supplementary Materials

### Figures S1 to S7

**Figure S1.**
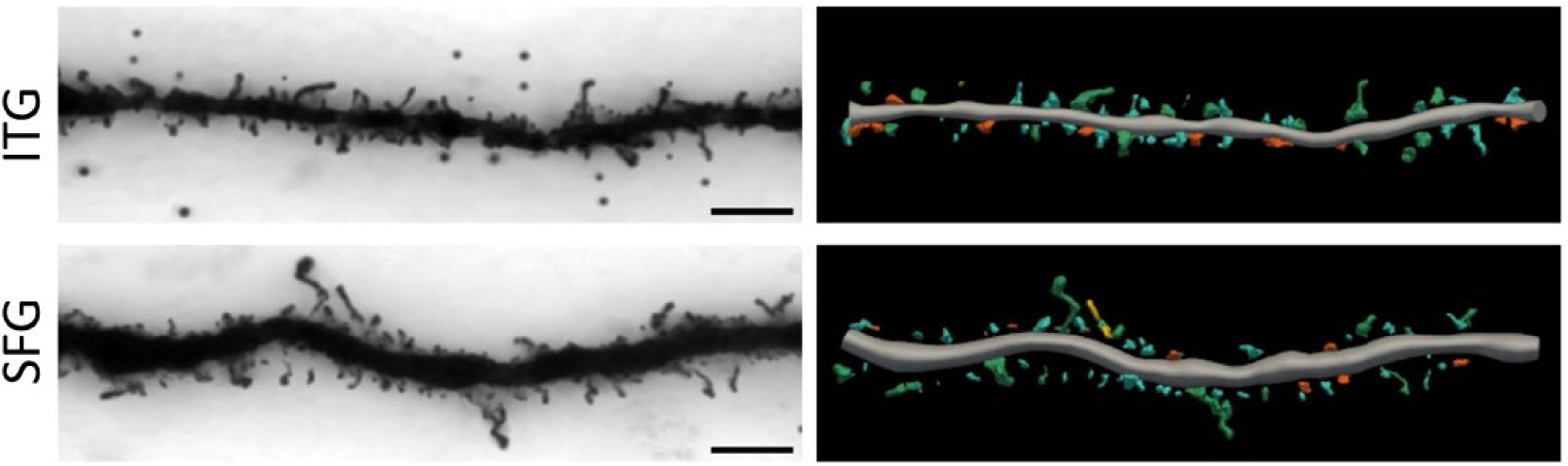
Representative microscopy images of dendrites and dendritic spines. 60X brightfield images of Golgi-stained dendrites from ITG (top) and SFG (bottom) of an exemplar subject are shown on the left. Scale bars = 10 µm. The digital three-dimensional reconstructions of those segments are shown on the right, which were used for estimating morphometric attributes.

**Figure S2.**
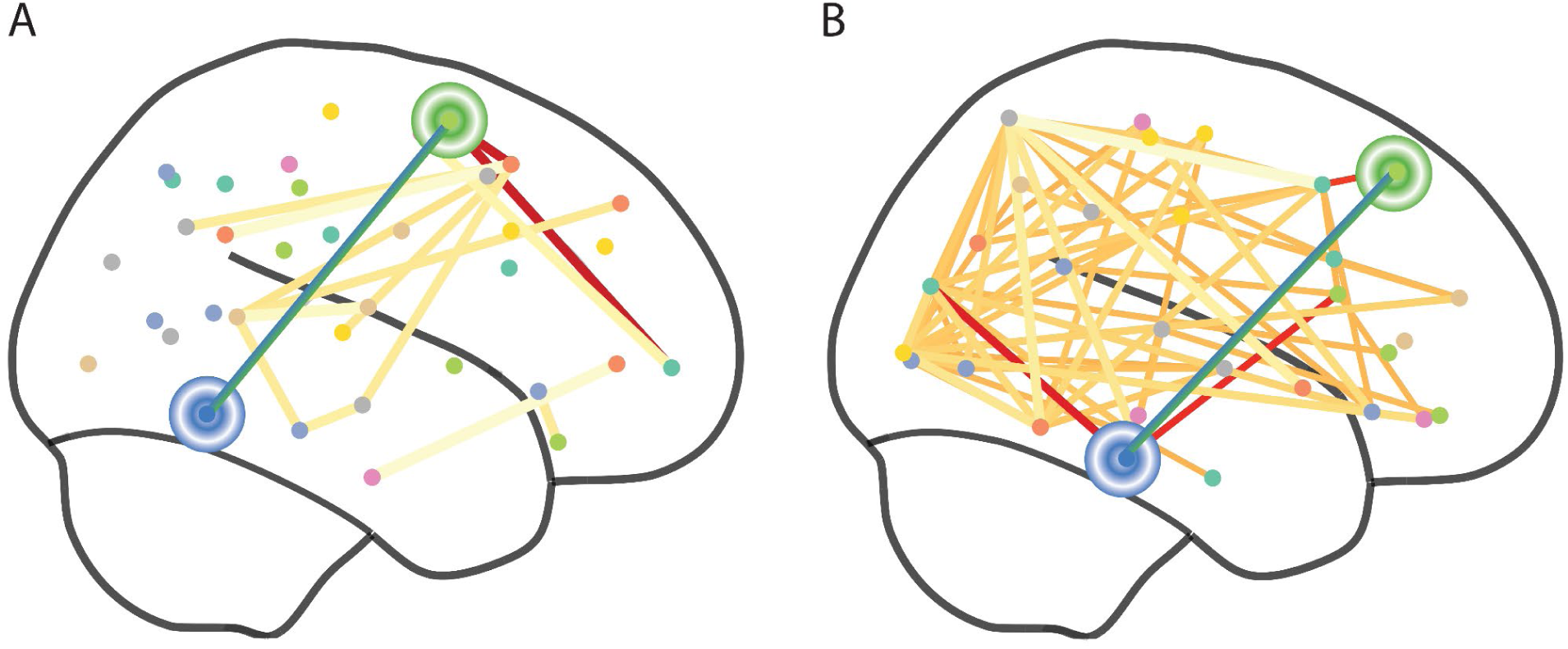
Association between brain attributes of interest and synaptic expression modules. A) Functional connectivity. Connections showing higher association strength with synaptic expression modules than the SFG-ITG connection displayed along with the SFG-ITG connection. Red corresponds to connections that emanate from either SFG or ITG, and yellow otherwise. Blue-green edge is between SFG and ITG. B) Structural connectivity.

**Figure S3.**
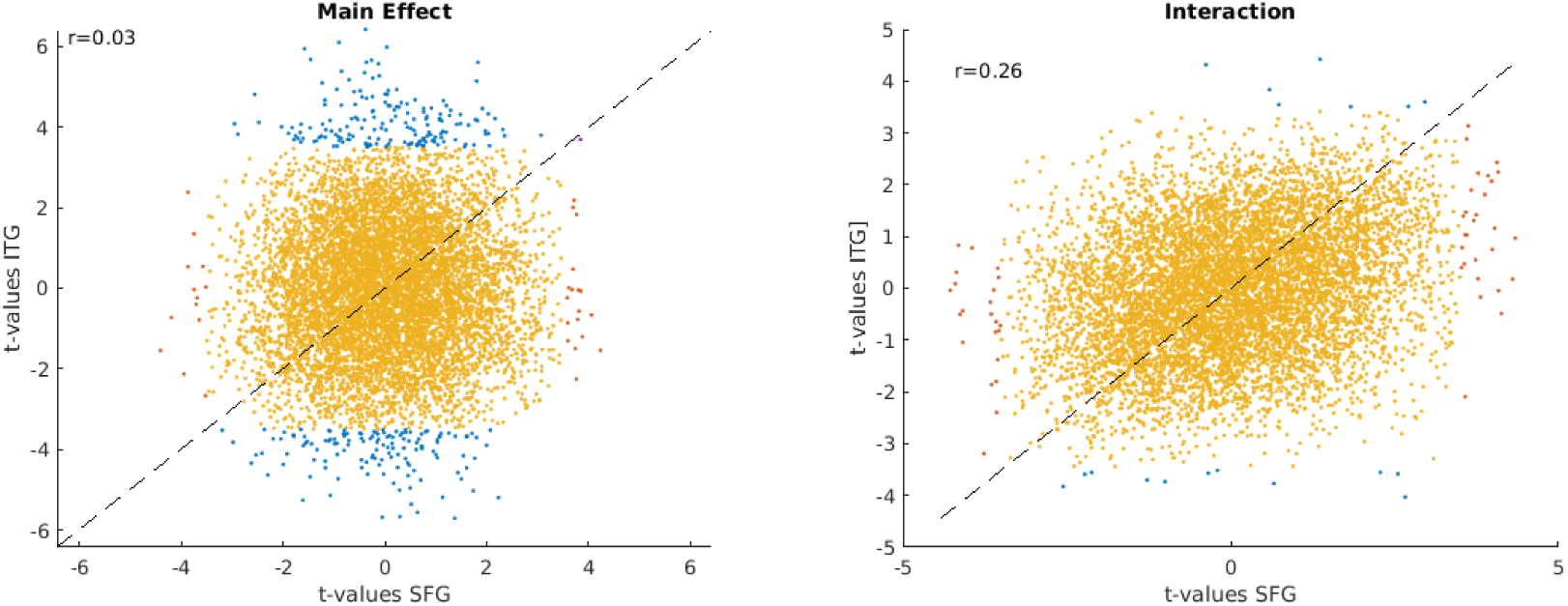
SFG vs. ITG proteins in association with functional connectivity. Each dot corresponds to a protein. Red and blue indicates significant for SFG only and ITG only, respectively. Purple indicates significant for both SFG and ITG.

**Figure S4.**
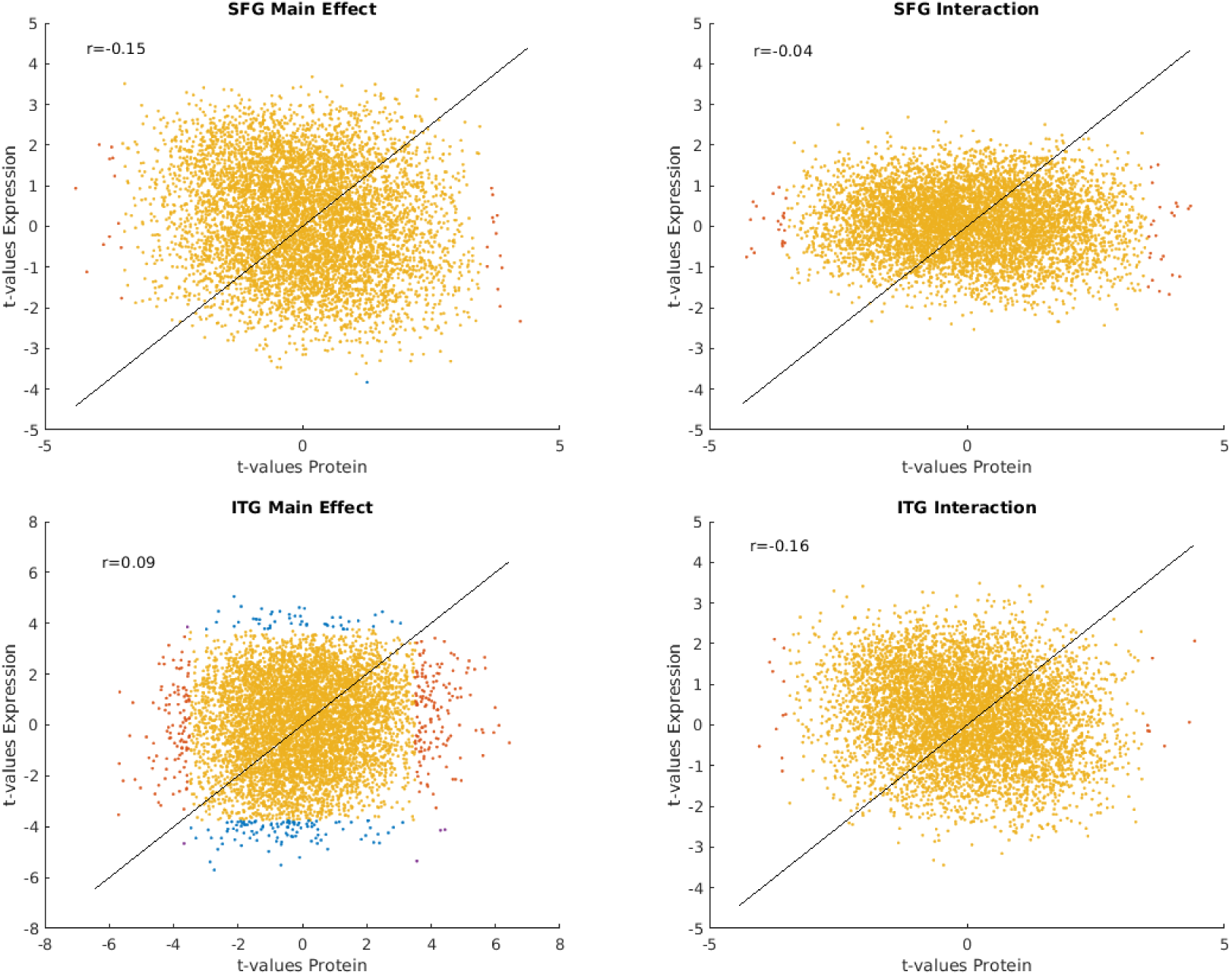
Protein abundance vs. gene expression in association with functional connectivity. Each dot corresponds to a gene/protein. Red and blue indicates significant for protein abundance only and gene expression only, respectively. Purple indicates significant for both protein abundance and gene expression.

**Figure S5.**
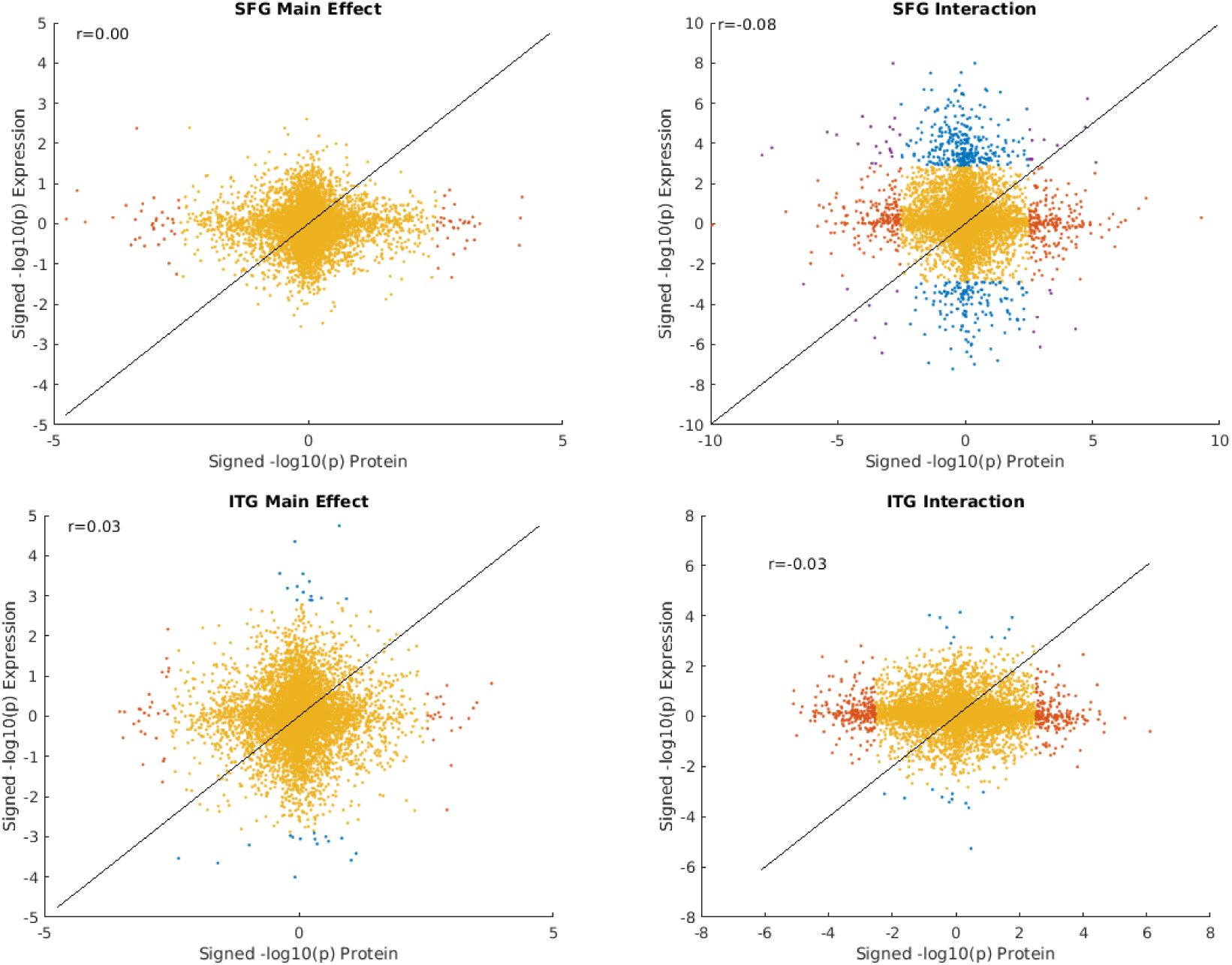
Protein abundance vs. gene expression in association with structural covariation. Each dot corresponds to a gene/protein. Red and blue indicates significant for protein abundance only and gene expression only, respectively. Purple indicates significant for both protein abundance and gene expression.

**Figure S6.**
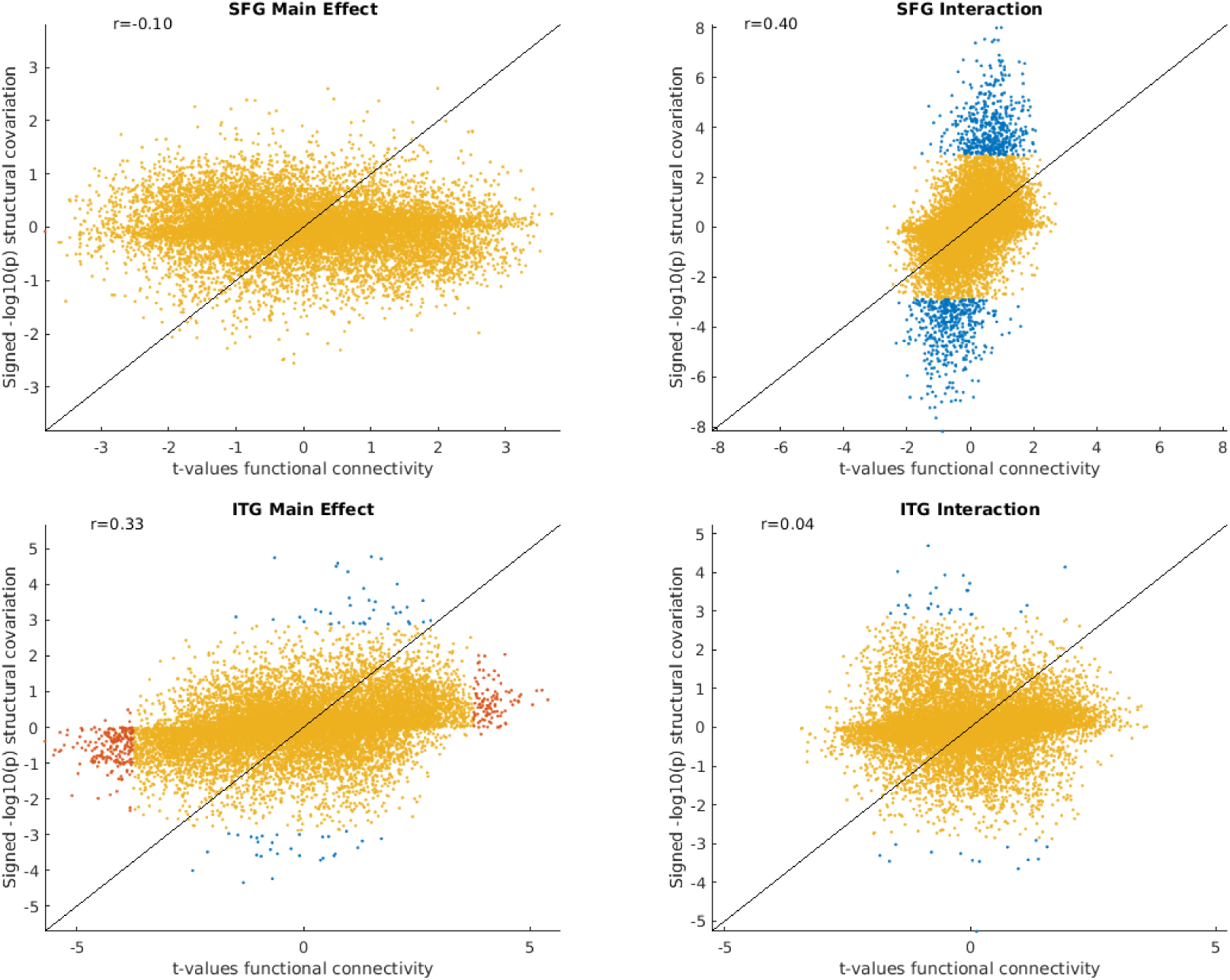
Functional connectivity vs. structural covariation in association with gene expression. Each dot corresponds to a gene. Red and blue indicates significant for functional connectivity only and structural covariation only, respectively. Purple indicates significant for both functional connectivity and structural covariation.

**Figure S7.**
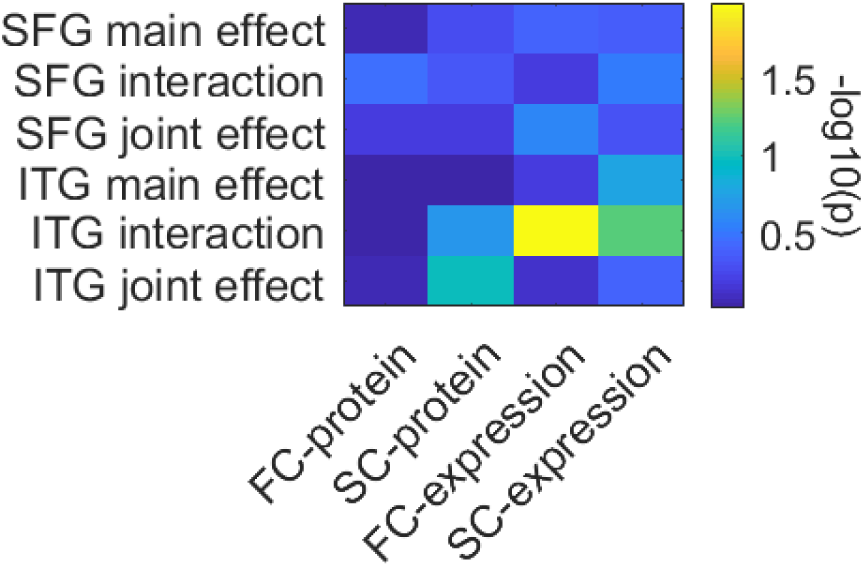
GSEA on 125 connectivity-related genes derived from ABHA. FC and SC denote functional connectivity and structural covariation, respectively.

### Tables S1 to S7

**Table S1. Protein abundance modules.** pModID indicates the module assignment of the SFG and ITG proteins. Only enriched GO terms with Bonferroni correction (p<5×10^-6^) listed.

**Table S2. Gene expression modules.** eModID indicates the module assignment of the SFG and ITG genes. Only enriched GO terms with Bonferroni correction (p<5×10^-6^) listed.

**Table S3. Summary statistics on associations between proteins and brain attributes.** FC and SC denote functional connectivity and structural covariation.

**Table S4. GO enrichment for proteins.** Enrichments were based on associations between the spine component of proteins and either functional connectivity (FC) or structural covariation (SC). Only enriched GO terms with Bonferroni correction (p<5×10^-6^) listed.

**Table S5. Summary statistics on associations between genes and brain attributes.** FC and SC denote functional connectivity and structural covariation.

**Table S6. GO enrichment for genes.** Enrichments were based on associations between the spine component of genes and either functional connectivity (FC) or structural covariation (SC). Only enriched GO terms with Bonferroni correction (p<5×10^-6^) listed. Note no GO terms passed p<5×10^-6^ for interaction effect on structural covariation for both SFG and ITG.

**Table S7. Protein level GWAS statistics.** MetaXcan was applied to convert SNP level GWAS z-scores to protein level z-scores.

